# Accelerated selection of a viral RNA polymerase variant during gene copy number amplification promotes rapid evolution of vaccinia virus

**DOI:** 10.1101/066845

**Authors:** Kelsey R. Cone, Zev N. Kronenberg, Mark Yandell, Nels C. Elde

**Author notes:** Present address: Department of Genome Sciences, University of Washington School of Medicine, Seattle, WA, USA.

## Abstract

Viruses are under relentless selective pressure from host immune defenses. To study how poxviruses adapt to innate immune detection pathways, we performed serial infections of vaccinia virus in primary human cells. Independent courses of experimental evolution with a recombinant strain lacking E3L revealed several high frequency point mutations in conserved poxvirus genes, suggesting important roles for essential poxvirus proteins in innate immune subversion. Two distinct mutations were identified in the viral RNA polymerase gene A24R, which seem to act through different mechanisms to increase virus replication. Specifically, a Leu18Phe substitution in A24R conferred fitness tradeoffs, including increased activation of the antiviral factor Protein kinase R (PKR). Intriguingly, this A24R variant underwent a drastic selective sweep during passaging, despite enhanced PKR activity. We show that the sweep of this variant can be accelerated by the presence of copy number variation (CNV) at the K3L locus, which with multiple copies strongly reduces PKR activation. Therefore, adaptive cases of CNV can also facilitate the accumulation of point mutations separate from the expanded locus. This study reveals how rapid bouts of gene copy number amplification during accrual of distant point mutations can potently facilitate poxvirus adaptation to host defenses.

**Importance:** Viruses can quickly evolve to defeat host immune functions. For poxviruses, little is known about how multiple adaptive mutations might concurrently emerge in populations. In this study, we uncovered a means of vaccinia virus adaptation involving the accumulation of distinct genetic variants within a single population. We identified adaptive point mutations in the viral RNA polymerase gene A24R, and show that these mutations can affect the activation of host nucleic acid sensing pathways through different mechanisms. We also found that structural variants within viral genomes in the form of gene copy number variation (CNV) provide dual benefits to evolving populations, including evidence that CNV facilitates the accumulation of a point mutation distant from the expanded locus. Our data suggest that transient CNV can accommodate the accumulation of mutations conferring modest benefits, or even fitness tradeoffs, and highlight how structural variation might aid poxvirus adaptation through both direct and indirect action.

## Introduction

Although the mutation rates of animal viruses are much higher than their hosts, the point mutation rate varies greatly between different types of viruses (1–4). For example, some double-stranded DNA (dsDNA) viruses have point mutation rates orders of magnitude lower than RNA viruses (3). Poxviruses, for instance, are predicted to have relatively low point mutation rates due to 3′–5′ proofreading activity of the viral DNA polymerase (3, 5, 6). While recent estimates suggest a higher point mutation rate for poxviruses compared to other dsDNA viruses (7, 8), these rates are still lower than that of most RNA viruses. Observations that viruses with varying mutation rates flourish in shared hosts strongly predict that successful adaptation of dsDNA viruses, including poxviruses, relies on mechanisms in addition to the rapid sampling of point mutations.

Vaccinia virus (VACV) provides a useful model system for studying poxvirus evolution due to the vast repertoire of available molecular tools and numerous well-characterized interactions between VACV-encoded factors and the host innate immune system. One key interface involves interactions between VACV and the host nucleic acid sensor Protein kinase R (PKR). Upon binding viral dsRNA, PKR phosphorylates the eukaryotic translation initiation factor eIF2α. Phosphorylation of eIF2α leads to a severe block in protein translation and attenuated viral replication. Like many poxviruses, VACV encodes E3L and K3L, which inhibit PKR via different mechanisms (9–11).

Consistent with their role as key host range factors, VACV E3L and K3L vary in their ability to block PKR from different host species (9, 12). In particular, K3L is a poor inhibitor of human PKR, such that a VACV strain lacking E3L (ΔE3L) (13) exhibits a severe replication defect during infection in HeLa cells (12). This deficiency places strong selective pressure on the virus to adapt to counteract PKR. Previous courses of experimental evolution of ΔE3L in HeLa cells revealed a recombination-based “genomic accordion” mechanism, in which copy number variation (CNV) of the K3L gene allowed rapid adaptation by inhibiting the activity of PKR (14). Although whole-genome sequencing of VACV revealed that each of the three replicate populations in this experiment acquired increased K3L copy number, there were differences between populations in the recombination breakpoints as well as unique high frequency point mutations throughout the genome. These differences suggest that experimental evolution is just beginning to uncover adaptive mutations and mechanisms contributing to poxvirus evolution. Further analysis of evolving populations under different conditions might reveal unrecognized means of virus adaptation.

In this study, we identified adaptive mutations in VACV genomes that arose during serial infections of primary human fibroblast (1°HF) cells. Interestingly, two nonsynonymous point mutations, from independent replicate populations, were identified within the A24R gene encoding a catalytic subunit of the viral RNA polymerase (vRNAP). Experimental rescue analysis indicates that either a Leu18Phe or Lys452Asn amino acid change in A24R is sufficient to provide a replication gain to the ΔE3L virus in 1°HF cells. Additionally, the A24R mutations we found seem to act through distinct mechanisms in comparison to previously identified adaptive A24R mutations to improve viral fitness. We also show that by blocking PKR activation, the K3L CNV arising in our virus populations enhanced the accumulation of a point mutation in A24R. This work provides a new view for how the selective sweep of a point mutation can occur concurrently with recombination-mediated adaptation in a viral population, and illuminates a fundamental mechanism for how structural variants might enhance poxvirus adaptation.

## Materials and Methods

### Cells and viruses

1°HF cells derived from human foreskin were a gift from Adam Geballe (Fred Hutchinson Cancer Research Center). 1°HF, HeLa, and BHK cells were maintained in Dulbecco’s modified Eagle’s medium (DMEM; HyClone) supplemented with 10% fetal bovine serum (HyClone), 1% penicillin-streptomycin (GE Lifesciences), and 1% stable L-glutamine (GE Lifesciences). Rhesus fibroblasts (Macaca mulatta, Coriell Institute for Medical Research #AG06252) and African green monkey fibroblasts (Cercopithecus aethiops, Coriell Institute for Medical Research #PR01193) were maintained in Minimum essential medium, alpha modification (MEM-alpha; HyClone) supplemented as above for DMEM. The Copenhagen strain of vaccinia virus (VC2) and the E3L deletion (ΔE3L) virus (13) were generous gifts from Bertram Jacobs (Arizona State University).

### Experimental evolution

For each infection during experimental evolution, 150mm dishes were seeded with an aliquot from the same stock of 1°HF cells (5x10^6^ cells/dish). Triplicate dishes of cells were infected (MOI = 1.0 for P1, MOI = 0.1 for subsequent passages) with ΔE3L virus for 2 hours in a minimal volume, and supplemented with media. At 48 hours, cells were collected, washed, pelleted, and resuspended in 1mL of media. Virus was released by one freeze/thaw cycle followed by sonication. Viral titers were determined by 48-hour plaque assay in BHK cells between each passage. Following 10 passages, an equal volume of virus from every other passage was expanded in BHK cells for 48 hours, and viral titers determined by 48-hour plaque assay in BHK cells, or 72 hours for replicate A passage 10 due to a small plaque phenotype.

### VACV whole-genome deep sequencing

Total viral genomic DNA was collected following 24-hour infection of BHK cells (MOI = 0.1) as previously described (15). Libraries were constructed using the Nextera XT DNA sample prep kit (Illumina, Inc.). Barcoded libraries were combined and sequenced using an Illumina MiSeq instrument at the High-Throughput Genomics Core (University of Utah). Reads (accession SRP073123) were mapped to the VC2 reference genome (accession M35027.1; modified on poxvirus.org) (16) using BWA mem (v0.7.10) (17) in default mode. PCR duplicates were removed and read depth was calculated using samtools (v0.1.18) (18). We utilized Genome Analysis Tool Kit (v3.2–2) (19) for base quality score recalibration, indel realignment, and variant calling across all samples (20, 21). We utilized Wham (v1.7.0–272–g078c-dirty) for structural variant calling (22). SNP and depth plots were generated in R (https://www.r-project.org/).

### Recombinant virus generation

500bp of homology surrounding the Leu18Phe-causing mutation in A24R was amplified by PCR from replicate A passage 10 viral DNA using primers A24R_1F: 5′ – C C T C C T C T C G A G C C C T C T C T G T T A G A T G A G G A T A G C and A24R_1R: 5′ – C C T C C T A C T A G T C A G T G A A C G T G G C T A A T G C G. 500bp of homology surrounding the Lys452Asn-causing mutation in A24R was amplified by PCR from VC2 viral DNA using primers A24R_2F: 5′ – C C T C C T C T C G A G C G T T G G C A C A T G A T G A A T T A G A G A A T T A C and A24R_2R: 5′ – C C T C C T A C T A G T G A G A T G C G A C T A G A G C A T T T T C T A T A G T G. The resulting PCR products were digested with XhoI and SpeI (New England Biolabs), gel purified, and cloned into pEQ1422 (gift from A. Geballe, Fred Hutchinson Cancer Research Center) (23) cut with the same enzymes to generate pEQ1422-Leu18Phe and pEQ1422-A24R_2. The Lys452Asn-causing mutation was introduced into pEQ1422-A24R_2 with site-directed mutagenesis primers A24R_Lys452Asn_F: 5′ – G T T G G A T T T T A T C C G G A T C A A G T A A A T A T T T C A A A G A T G T T T T C T G T C A and A24R_Lys452Asn_R: 5′ – T G A C A G A A A A C A T C T T T G A A A T A T T T A C T T G A T C C G G A T A A A A T C C A A C (pEQ1422-Lys452Asn). BHK cells were infected with ΔE3L (MOI = 1.0), and transfected 1 hour post-infection with pEQ1422-Leu18Phe or pEQ1422-Lys452Asn using FuGENE6 (Promega) according to the manufacturer’s protocol. Infected cells were collected 48 hours post-infection, and viruses released by one freeze/thaw cycle followed by sonication. Resulting viruses were selected using transient dominant selection (24). Briefly, viruses were plaque purified three times in the presence of 600μg/mL hygromycin B (Sigma-Aldrich), followed by three rounds of plaque purification without hygromycin B. The presence of the Leu18Phe or Lys452Asn substitution and loss of HygR were confirmed by PCR followed by Sanger sequencing. Viruses were amplified and titered in BHK cells.

### One-step growth curve

BHK or 1°HF cells were infected with ΔE3L or A24R^Leu18Phe^ VACV (MOI 2.5) in duplicate, and virus was replaced with fresh media after 2 hours. Cells were harvested at 2, 6, 12, 24, 48, and 72 hours post infection, and viral titers determined by 72-hour plaque assay in BHK cells.

### Modeling

Amino acid alignment between VACV A24R and *S. cerevisiae* RBP2 was generated using Clustal Omega (v1.2.1). Corresponding A24R variant residues were then mapped onto the *S. cerevisiae* RNAP II crystal structure (PDB: 1I50) (25) using Chimera software (http://www.cgl.ucsf.edu/chimera/) (26).

### rRNA degradation assay

1°HF cells were pretreated with interferon-alpha for 24 hours. Cells were then left untreated (mock), transfected with Poly(I:C), or infected with ΔE3L, A24R^Leu18Phe^, or A24R^Lys452Asn^ virus (MOI 5.0) for 6 hours. Total RNA was harvested and 225ng/lane was run on an Agilent 2200 tapestation.

### Immunoblot analysis

1°HF cells were untreated (mock), or infected with ΔE3L, A24R^Leu18Phe^, A24R^Lys452Asn^, or replicate A P10 virus (MOI 5.0) for 6 hours. Protein lysates were collected in RIPA lysis buffer, and total protein concentration was quantified by Bradford assay using a Synergy HT plate reader (BioTek). Equivalent amounts of lysate were separated on a pre-cast Mini-PROTEAN TGX gel (Bio-Rad), and transferred to a PVDF membrane (Immobilon). Blots were blocked for 30 minutes in PBST + 5% milk, or for 10 minutes followed by three washes for phospho antibodies. Blots were then incubated with the following primary antibody overnight at 4°C: MαPKR B-10 (Santa Cruz; 1:200), RαPhospho-PKR E120 (Abcam; 1:500), RαeIF2α (Cell Signaling; 1:1000), MαPhospho-eIF2α (Cell Signaling; 1:250). Blots were probed with the appropriate secondary antibody for 1 hour at RT: GαM-IgG-HRP (Millipore; 1:50,000), GαR-IgG-HRP (Miillipore; 1:50,000), activated with WesternBright ECL reagent (Advansta), and exposed to autoradiography film (GeneMate) developed in a Mini-Med 90 film processor (AFP Imaging).

### Plaque size analysis

BHK cells were infected with ΔE3L or A24R^Leu18Phe^ virus (MOI 0.1) for 48 hours, and stained with crystal violet. Plates were imaged on a Gel Doc^TM^ XR+ system (BioRad), and plaque size quantified using ImageJ v1.48 software (Rasband). Three independent wells for a total of 350 plaques were analyzed for each virus.

### Southern blot analysis

Total viral DNA was collected as above, and ~2μg per sample was digested with BspEI (New England Biolabs). Digested DNA was separated by agarose gel electrophoresis, and transferred to nylon membranes (GE Lifesciences) using a vacuum transfer, followed by UV-crosslinking. The resulting blots were probed with PCR-amplified K3L using the DIG High-Prime DNA Labeling & Detection Starter Kit II (Roche) according to the manufacturer’s protocol.

### Mutation accumulation assay

Replicate A P10 virus was passaged an additional three times in 1°HF cells until the A24R Leu18Phe variant was fixed (A24R^Leu18Phe^+CNV virus). The K3L CNV alone virus contains the same CNV breakpoint, but has no mutations in A24R (CNV virus). BHK cells were infected with viruses mixed at a ratio of 1:100 as follows: A24R^Leu18Phe^:ΔE3L, CNV:ΔE3L, or A24R^Leu18Phe^+CNV:ΔE3L for two passages in BHK cells (MOI 0.1). All passages were collected after 48 hours as above, and titered by 72-hour plaque assay in BHK cells.

### Deep sequencing data

All deep sequencing data are available on the Sequence Read Archive, under accession SRP073123.

## Results

### Replication gains following serial infection of 1°HF cells

To study mechanisms of viral adaptation, we performed serial infections of VACV in primary human fibroblast (1°HF) cells. ΔE3L replicates poorly in HeLa cells (12), so we first tested whether the defect is similar for infections in primary human fibroblasts. While both ΔE3L and the parental wild-type Copenhagen strain of VACV (VC2) replicate equally in a permissive hamster (BHK) cell line during a 48 hour infection, ΔE3L displays an even stronger growth defect in 1°HF cells than in HeLa cells when compared to VC2 (Figure 1A). This result is consistent with cell type-specific innate immune responses to viral infection placing different selective pressure on the virus (27), even between different human cell lines.

**Figure 1.**
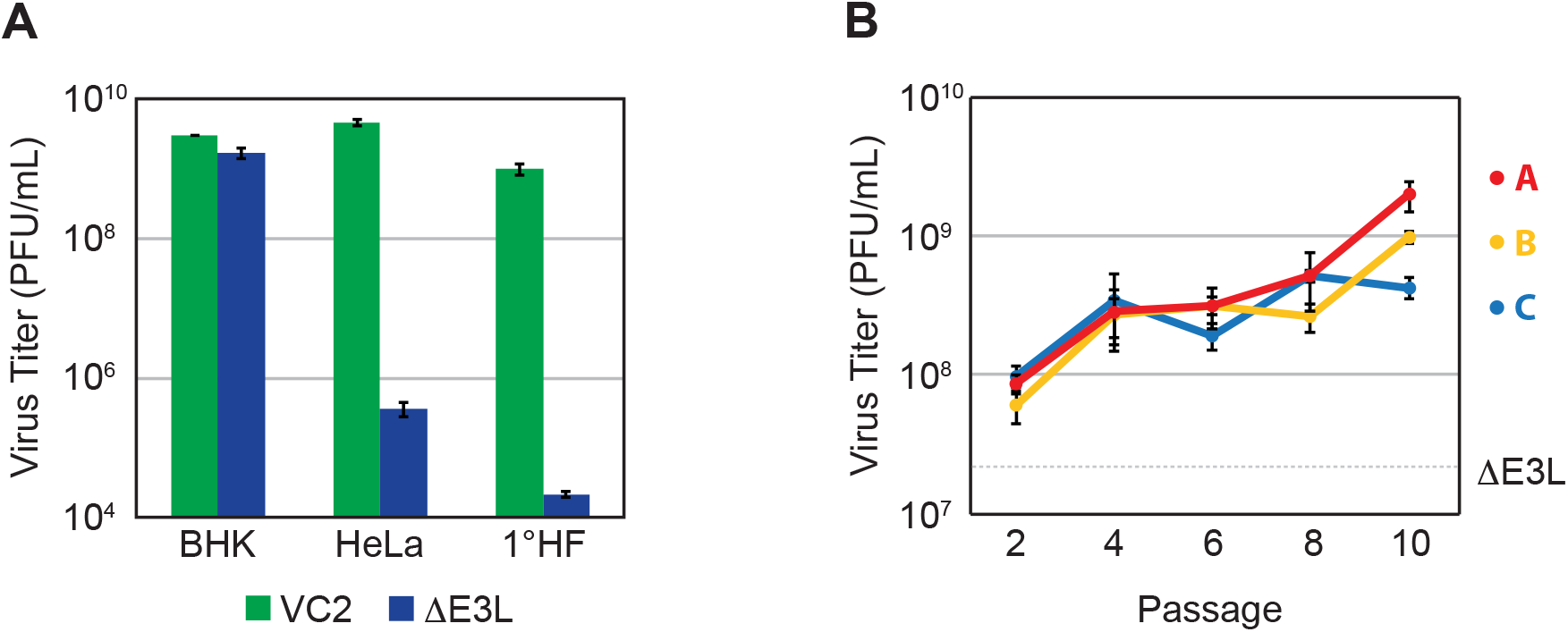
Rapid adaptation of ΔE3L during experimental evolution in 1°HF cells. (A) Cells were infected with wild-type VC2 or ΔE3L virus (MOI 0.1) for 48 hours. (B) Triplicate populations of ΔE3L virus were passaged 10 times in 1°HF cells. Vreus populationt from every other passage were expanded in BHK cells, and titered simultaneously. Input parental ΔE3L is indicated by the dotted line. All viral titers were measured in BHK cells by 48-hour plaque assay, as mean PFU/mL ± standard deviation.

To test how VACV might adapt to primary human cells, we performed serial infections of ΔE3L in triplicate in 1°HF cells. Using viral titer as a measure of fitness, we observed rapid gains in fitness over the course of ten passages in all three replicate virus populations (Figure 1B). Despite modest replication of the parental ΔE3L virus in 1°HFs compared to HeLa cells, we observed comparable gains in replication of ΔE3L virus populations over the course of our infections in 1°HF cells (Figure 1B), and HeLa cells as previously reported (14).

### High frequency point mutations in evolved virus populations

To define genetic changes that might account for increases in viral fitness, we used deep sequencing of viral genomes from each of the replicate populations after ten rounds of serial infection (P10). We obtained an average of >2,000x coverage across the genome for each P10 population. Excluding the inverted terminal repeat regions (1–5,000bp, 186,737–191,737bp), we identified nine single nucleotide polymorphisms (SNPs) not present in the ΔE3L parent virus at a frequency greater than 1% in any P10 population, in addition to twenty shared differences compared to the VC2 reference strain (Table 1). The nine SNPs present in at least one of the three P10 populations but not in the ΔE3L parent virus represent potentially adaptive mutations. Notably, all nine SNPs lie within open reading frames, and seven of these resulted in a nonsynonymous amino acid change or frameshift (Figure 2A). Remarkably, the highest frequency SNP in each population caused a substitution in an essential gene conserved among poxviruses: a Leu18Phe amino acid substitution in A24R, a catalytic subunit of the viral RNA polymerase (vRNAP) (28); a Glu495Gly amino acid substitution in E9L, the viral DNA polymerase (29, 30); and a Trp44Cys amino acid substitution in F10L, a kinase required for virion morphogenesis (31). Furthermore, a mutation causing a Lys452Asn amino acid substitution in A24R was identified in an independent replicate population from the A24R Leu18Phe variant. This result suggests that the A24R gene may be a common target for beneficial mutations in VACV, consistent with previous reports of other adaptive A24R mutations (23, 32, 33). Together these high frequency mutations suggest a common role for poxvirus genes encoding essential viral functions in the adaptation to host innate immune responses.

**Figure 2.**
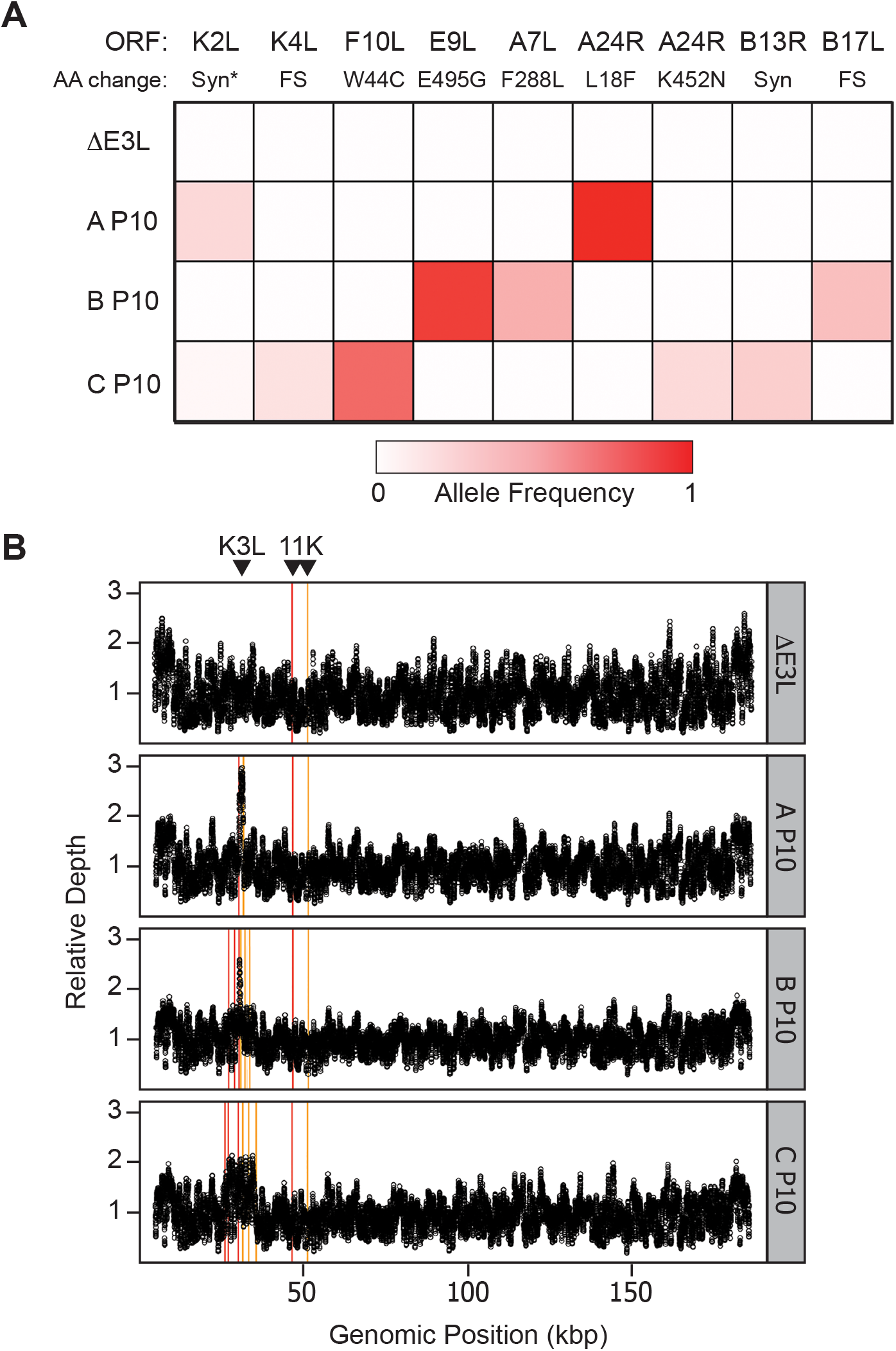
Genetic changes in virus populations following experimental evolution. (A) Allele frequencies were obtained by deep sequencing of viral genomic DNA from passage 10 virus populations compared to ΔE3L parent virus in 1°HF cells. Alleles with a frequency ≥0.01 not present in the parent virus are shown. The VACV open-reading frame (ORF) and resulting amino acid change are listed above. FS = frameshift, Syn = synonymous. *The K2L allele is likely a miscalled structural variant. (B) Relative depth of coverage across the viral genome is shown in black, excluding the inverted terminal repeat regions. Breakpoint positions of structural variants called by Wham are shown as vertical lines, with 5′ positions in red and 3′ positions in orange. Genomic location of the K3L gene and the duplicated 11K promoter are indicated above.

**Table 1.**
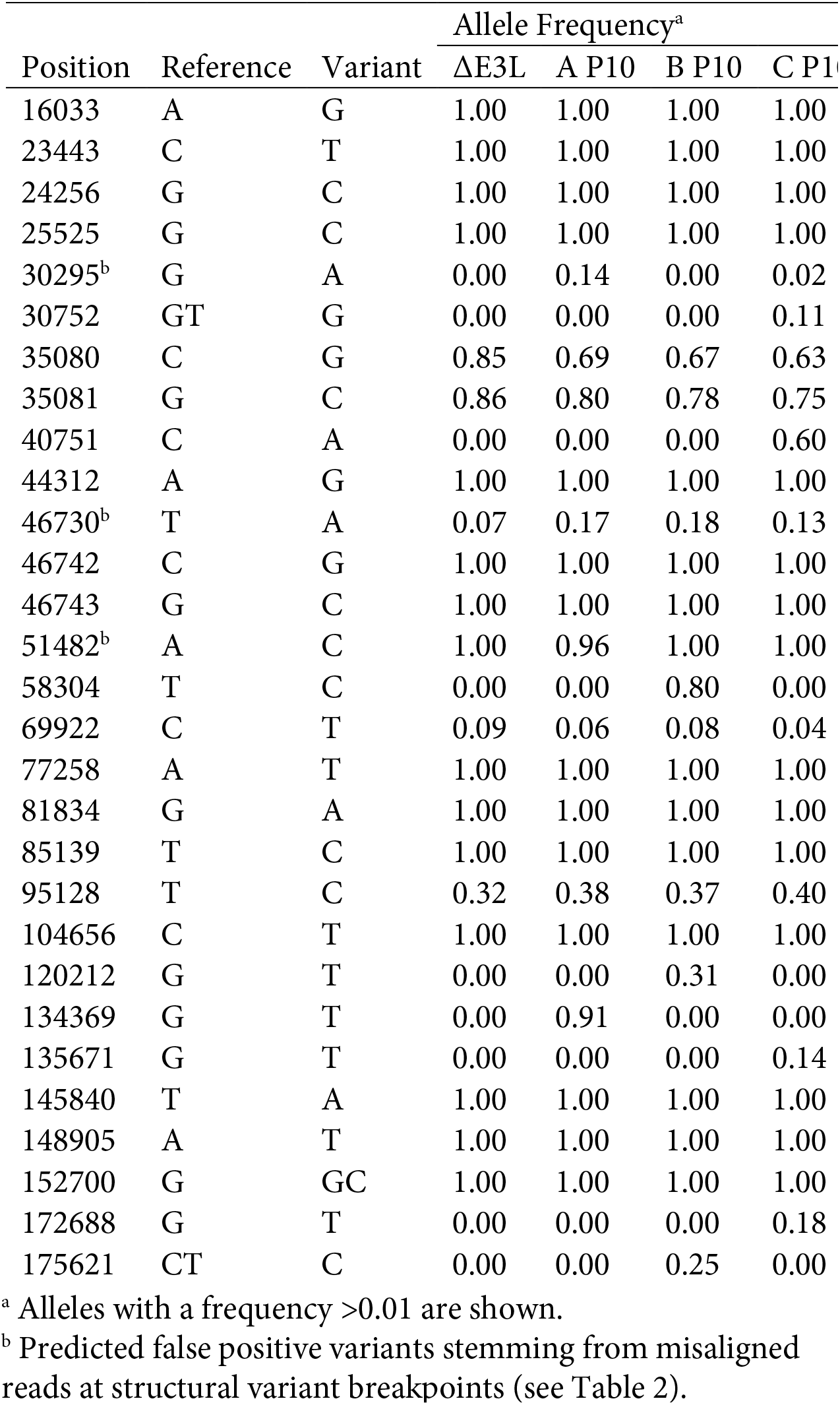
Point mutations following passaging in 1°HF cells

### Structural variants in evolved virus populations

In addition to point mutations, genetic changes in the form of gene copy number variation (CNV) have previously been shown to play an adaptive role during poxvirus adaptation (14). To identify potentially adaptive structural variants (SVs) in our evolved virus populations, we analyzed the P10 sequences using SV analysis implemented in the program Wham (22). We found seven SVs with >10 reads to define both the 5′ and 3′ location of recombination breakpoints in the virus populations (Figure 2B, Table 2). Sanger sequencing corroborated each of these SVs to within one nucleotide of the putative breakpoint. The single SV identified in the ΔE3L parent virus is also present in all three P10 populations (breakpoint 7 in Table 2). This variant corresponds to the 11K vaccinia promoter introduced during the deletion of E3L (13), resulting in two copies of this sequence at different genomic locations (Figure 2B). The remaining 6 SVs are only present in the P10 virus populations, and each of them is associated with the K3L locus. There is a corresponding increase in sequencing depth surrounding K3L in each P10 virus population but absent in the parent population (Figure 2B). Three of the SNPs from variant calling are located one base pair from a structural variant breakpoint, and we therefore categorized these as false positive calls due to misaligned reads (Table 1). This finding illustrates how SVs should be considered in poxvirus genomes when using standard variant calling methods to identify SNPs.

**Table 2.** Structural variants following passaging in 1°HF cells

| Breakpoint | Genomic Position<sup>a</sup> |  | Read Support (5', 3') |  |  |  |
| --- | --- | --- | --- | --- | --- | --- |
| | 5' | 3' | ΔE3L | A P10 | B P10 | C P10 |
| 1 | 26302 | 35775 | 0, 0 | 0, 0 | 0, 0 | 112, 102 |
| 2 | 27105 | 32160 | 0, 0 | 0, 0 | 74, 70 | 0, 0 |
| 3 | 27322 | 33524 | 0, 0 | 0, 0 | 0, 0 | 105, 73 |
| 4 | 28875 | 33568 | 0, 0 | 0, 0 | 114, 74 | 0, 0 |
| 5 | 30287 | 30840 | 0, 0 | 0, 0 | 164, 125 | 0, 0 |
| 6 | 30296 | 31725 | 0, 0 | 190, 137 | 0, 0 | 96, 85 |
| 7 | 46731 | 51483 | 179, 260 | 159, 465 | 179, 472 | 200, 389 |

<sup>a</sup> Positions supported by >10 reads are shown.

A similar copy number amplification of the K3L locus was observed following serial infections of ΔE3L in HeLa cells (14). In this previous study we showed that CNV at the K3L locus corresponds to increased K3L protein, which impairs PKR activation. Interestingly, two of the seven breakpoints identified in this study were identical to those observed in virus populations passaged in HeLa cells (14), suggesting the presence of these variants at a level below detection in the ΔE3L parent virus, or that these are preferential sites for recombination. These results, combined with studies demonstrating CNV in other poxvirus genes in response to selective pressure (34–36), continue to reveal CNV as a mechanism for rapid adaptation of VACV.

### A24R mutations increase viral fitness through distinct mechanisms

Following 10 passages in 1°HF cells, each population of viruses harbored a unique set of mutations that might contribute to increased fitness. The most drastic example was a Leu18Phe amino acid substitution in A24R that was nearly fixed (frequency = 0.91) in one virus population after 10 passages (Figure 2A). The lack of other nonsynonymous point mutations in this replicate population above a frequency of 1% strongly suggests that this single mutation contributes to the observed increase in fitness (Figure 1B). To test this hypothesis, we generated a recombinant virus with this A24R mutation in the parental ΔE3L strain (A24R^Leu18Phe^). Growth curve analysis in the permissive BHK cell line revealed that A24R^Leu18Phe^ and ΔE3L replicate to similar titers at 72 hours post-infection, even though the A24R^Leu18Phe^ virus lags behind ΔE3L at both 24 and 48 hours (Figure 3A). However, in 1°HF the A24R^Leu18Phe^ virus exhibits a significant increase in titer relative to ΔE3L from 24 to 72 hours post-infection. Thus, the single nucleotide change in A24R in a ΔE3L genetic background is sufficient to enhance viral fitness under selective pressure in 1°HF cells. To examine how a single amino acid substitution in A24R might increase viral fitness, we mapped A24R mutations onto solved structures of RNA polymerases. Previous work used the crystal structure of *S. cerevisiae* RNA polymerase II to map A24R amino acid substitutions onto the homologous subunit in yeast, RBP2 (33). RBP2 is the second largest subunit of yeast RNA polymerase II and forms an active site of the enzyme with RBP1 (37). Using a similar approach, we generated an amino acid alignment between VACV A24R and *S. cerevisiae* RBP2 to predict the location of A24R amino acid substitutions on the *S. cerevisiae* RNA polymerase II crystal structure (PDB: 1I50) (25). This analysis suggests that the A24R Leu18Phe substitution is located on a solvent-exposed surface of the polymerase distal from the active site of the enzyme (Figure 3B). This surface might be involved in binding other factors to the polymerase, such that a Leu18Phe substitution could alter protein-protein interaction(s). However, little is currently known about viral or host proteins that bind to A24R, making it difficult to predict functional consequences of this mutation.

**Figure 3.**
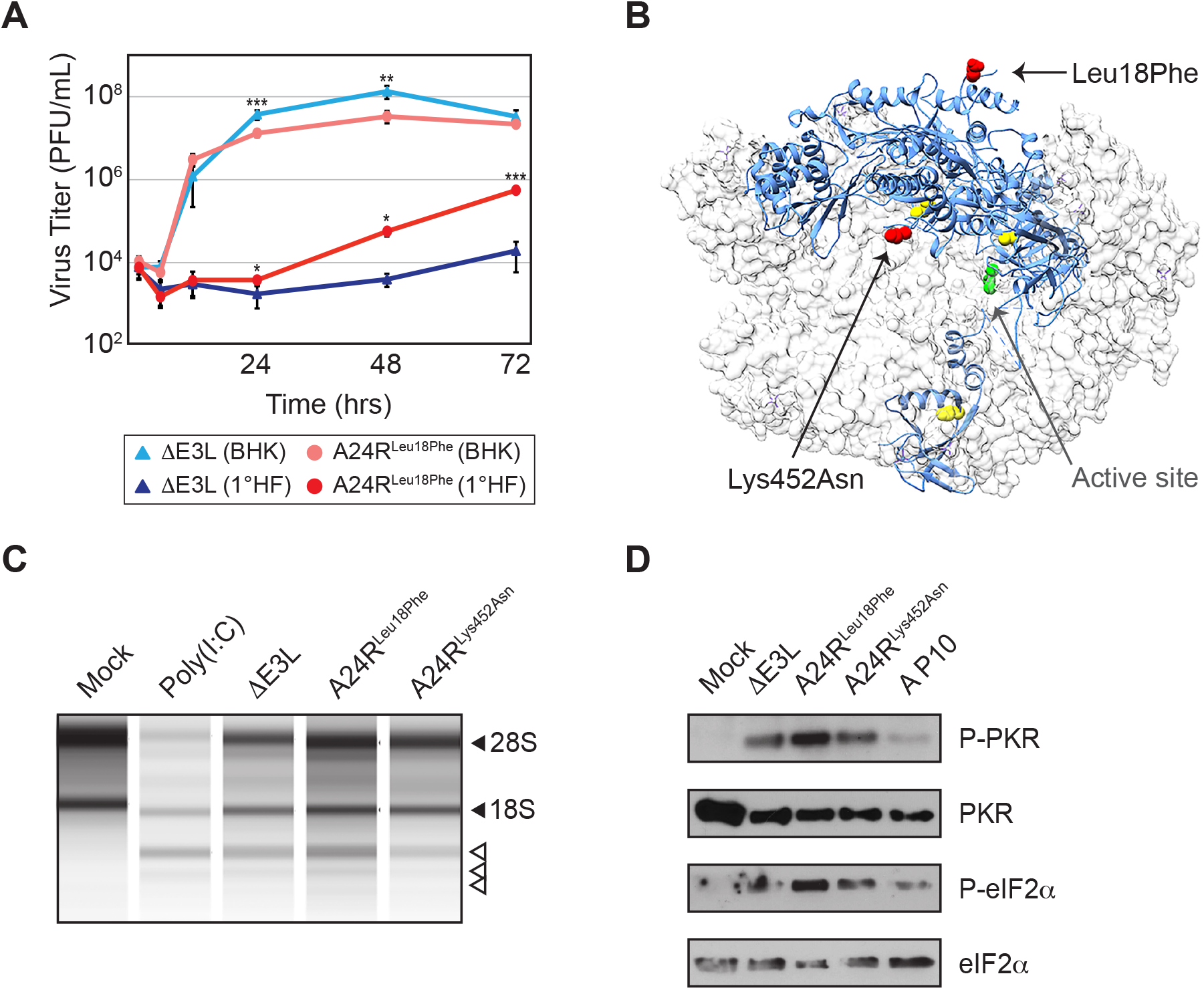
A24R mutations increase fitness through distinct mechanisms. (A) Single-step growth curve analysis was performed with either the ΔE3L or A24R^Leul8Phe^ recombinant virus in BHK or 1°HF cells. Viral titers were measured in BHK cells by 72-hour plaque assay, as mean PFU/mL ± standard deviation. *p<0.05, **p<0.01, ***p<0.005 by 2-tailed T-test relative to ΔE3L within each cell type. (B) Positions of A24R mutations identified in this study (red spheres) or previously published data (yellow spheres) were mapped onto the homologous *S. cerevisiae* RNAP II structure (PDB: 1150). The RBP2 protein (homologous to A24R) is shown in blue ribbons in the context of the multi-subunit polymerase in gray (active site in green). (C) rRNA degradation in 1°HF cells left untreated (mock), transfected with Poly(I:C), or infected with ΔE3L, A24R^Leul8Phe^, or A24R^Lys452Asn^ virus (MOI 5.0) for 6 hours. Filled arrowheads indicate 28S and 18S rRNA, and open arrowheads indicate degradation products. (D) Immunoblot for phosphorylated or total PKR and phosphorylated or total eIF2α in 1°HF cells infected with ΔE3L, A24R^Leu18Phe^, A24R^Lys452Asn^, or replicate A passage 10 virus (MOI 5.0) for 6 hours. A representative image of five independent blots is shown.

Distinct adaptive mutations in A24R demonstrate how vRNAP variation can impact viral fitness. Two A24R mutations were shown to influence transcription elongation, resulting in the production of short virus transcripts in response to drug selection (32, 33). These mutations also reduced the activation of the host dsRNA sensor oligoadenylate synthetase (OAS). These studies suggest that changes to transcript length, dictated by vRNAP, can influence the activation of innate immune dsRNA sensors in the cell. Indeed, another A24R mutation was recently identified that reduces the activation of the dsRNA sensor PKR (23). Without E3L, viruses are more vulnerable to PKR and OAS activity, and thus we hypothesized that the A24R substitutions we identified might act through a similar mechanism to reduce the activation of dsRNA sensors.

To test whether either of the A24R variants from our evolved virus populations is sufficient to alter innate immune pathways, we measured the activity of the OAS/RNaseL and PKR pathways in cells infected with recombinant viruses harboring either the Leu18Phe or Lys452Asn A24R substitutions. We did not detect any notable difference in RNaseL activity as judged by ribosomal RNA degradation following infection with either A24R^Leu18Phe^, or a similarly generated A24R^Lys452Asn^ recombinant virus containing the Lys452Asn substitution in A24R (Figure 3C). This result suggests that the OAS/RNaseL pathway is not affected by these A24R substitutions. We next tested whether either of the A24R variants reduce PKR activation, measured by changes in phosphorylated PKR and phosphorylated eIF2α protein levels in infected 1°HF cells. Counter to our expectation, immunoblot analysis indicated that 1°HF cells infected with the A24R^Leu18Phe^ virus repeatedly showed increased levels of both phosphorylated PKR and eIF2α (Figure 3D). While the A24R^Leu18Phe^ virus confers a replication benefit that could account for an increase in PKR activation, this is unlikely given equivalent viral titers between the A24R^Leu18Phe^ and ΔE3L viruses at the 6-hour timepoint (Figure 3A) when we harvested total protein. In contrast, there were no substantial changes in PKR activation upon infection with the A24R^Lys452Asn^ virus compared to infection with ΔE3L. These data suggest that the A24R variants we identified work through distinct mechanisms compared to other known A24R substitutions to enhance viral fitness. Moreover, the increase in PKR activation with the A24R^Leu18Phe^ virus was paradoxical, considering the replication advantage we observed in 1°HF cells.

### The A24R Leu18Phe variant confers fitness tradeoffs

Given that E3L and K3L act as host range factors, blocking innate immune activation in some hosts but not others (12), we tested the impact of A24R variation during infections of cells from other primate species. We performed 48-hour infections in two human and two old-world primate cell lines in duplicate: 1°HF, HeLa, rhesus macaque fibroblasts, and African green monkey fibroblasts. In each of the four cell lines tested, the parental ΔE3L virus exhibits considerably reduced replication compared to a permissive BHK cell line (Figure 4A). The A24R^Lys452Asn^ virus replicated equal to or better than ΔE3L in all primate cells tested, suggesting that the fitness increase provided by the A24R Lys452Asn substitution is not species- or cell type-specific. In contrast, while A24R^Leu18Phe^ virus replication showed a 25-fold increase in 1°HF and a smaller increase in rhesus macaque fibroblasts relative to ΔE3L virus, titers were reduced in HeLa cells and African green monkey fibroblasts. The clear differences in A24R^Leu18Phe^ viral titers between two Old World monkey species, and also between two human cell lines reveals a potential fitness tradeoff for viruses harboring this substitution.

**Figure 4.**
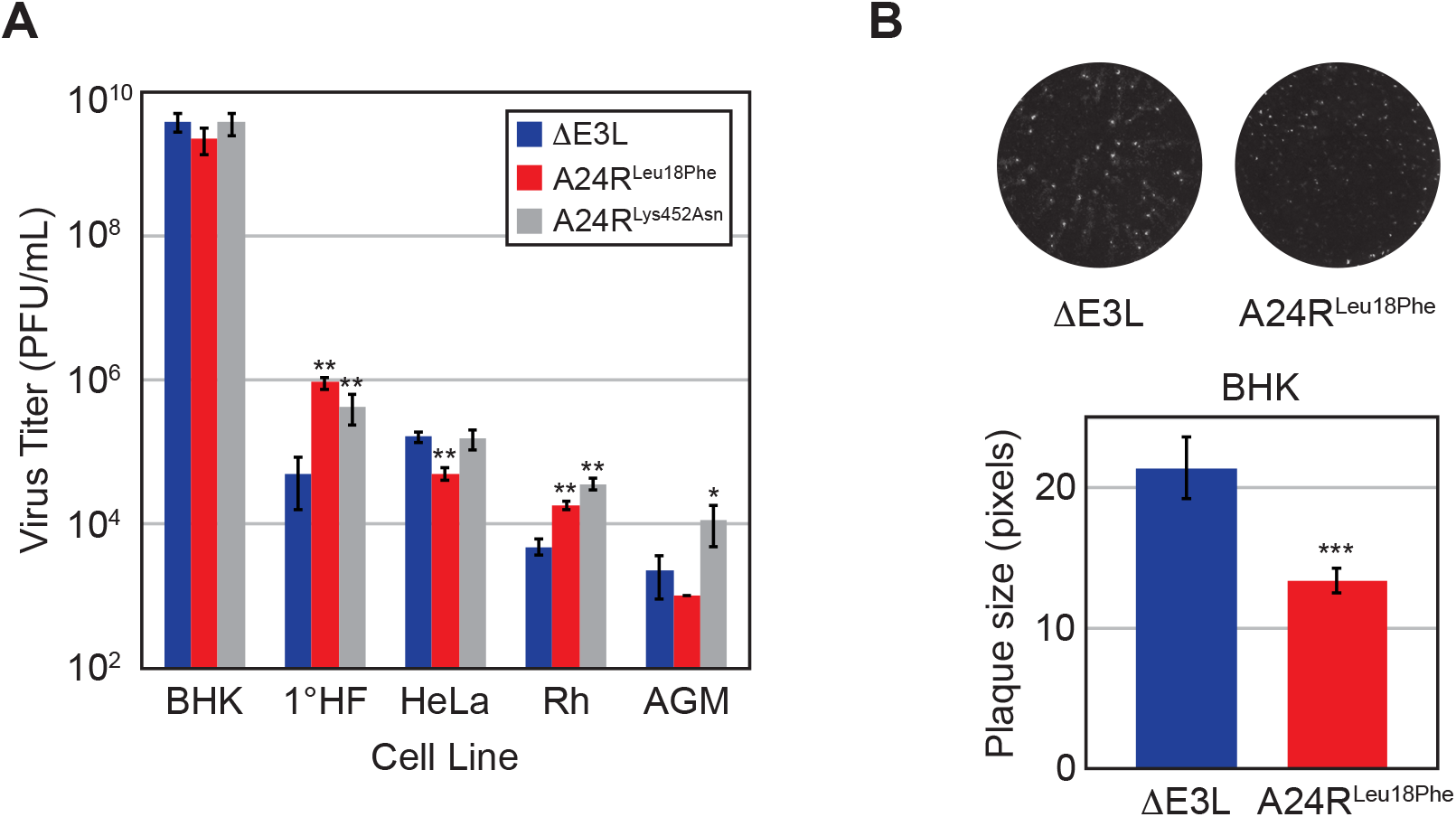
The A24R Leu18Phe variant confers fitness tradeoffs. (A) Cell lines were infected in duplicate with ΔE3L, A24R^Leul8Phe^, or A24R^Lys452Asn^ virus (MOI 0.1) for 48 hours. Viral titers were measured in BHK cells by 72-hour plaque assay, as mean PFU/mL ± standard deviation, *p<0.05, **p<0.01 by 2-tailed T-test relative to ΔE3L within each cell type. Rh = Rhesus. (B) Average plaque size ± standard deviation from three independent wells of BHK cells infected with ΔE3L or A24R^Leul8Phe^ virus (MOI 0.1) for 48 hours. A representative image is shown, ***p<0.005 by 2-tailed T-test.

Consistent with a fitness tradeoff in some cell types, the A24R^Leu18Phe^ virus displays a small plaque phenotype in permissive BHK cells (Figure 4B). This phenotype is associated with defects in cell-to-cell spread (38–40), and while the A24R^Leu18Phe^ virus produces approximately the same number of infectious particles as ΔE3L in BHK cells after 72 hours, the A24R^Leu18Phe^ virus lags behind ΔE3L at 24 and 48 hours (Figure 3A). The small plaque phenotype in conjunction with differential replication among cell lines supports the idea that the A24R Leu18Phe variant is beneficial only under certain conditions. Furthermore, the growth benefit in 1°HF cells and growth defect in HeLa cells highlights that specific, species-independent differences between the cell lines can contribute to the success or failure of the A24R^Leu18Phe^ virus.

### Accelerated sweep of the A24R Leu18Phe variant during adaptive gene copy number amplification

The A24R Leu18Phe variant rose to near fixation in a viral population during experimental evolution of ΔE3L in 1°HF cells (Figure 2, Table 1). The rapid dominance of the variant in this population was unusual, given the slower accumulation dynamics of mutations within other replicate populations. For example, the A24R Lys452Asn variant provides a similar replication increase alone (Figure 4A), yet only reaches a frequency of 14% compared to the A24R Leu18Phe variant at 91% (Figure 5A). Furthermore, we might have predicted that the increased PKR activation induced by the A24R Leu18Phe variant (Figure 3D) would prevent the mutation from reaching fixation so rapidly. To more carefully determine how this mutation arose, we analyzed earlier passages during the course of experimental evolution. Remarkably, the A24R Leu18Phe variant was only detectable starting at passage 8 as judged by sequencing A24R amplicons in each passage, prompting us to deep sequence virus populations from passages 7–9. Genome sequence analysis revealed that the mutation rapidly increased in frequency with each successive passage late in the experiment (Figure 5A). Consistent with our earlier analysis, this was the only verified SNP above 1% in passages 7–10 (replicate A), which is in stark contrast to replicate C in which multiple mutations across the genome fluctuated in frequency from passages 7–10 (Figure 5A). The rapid accumulation of the A24R Leu18Phe variant, as well as a lack of any other mutations across the genome, is consistent with a strong selective sweep of the mutation in the replicate A population. However, because the A24R^Leu18Phe^ virus also increased PKR activation, the question remained as to how the mutation induces increased virus replication.

**Figure 5.**
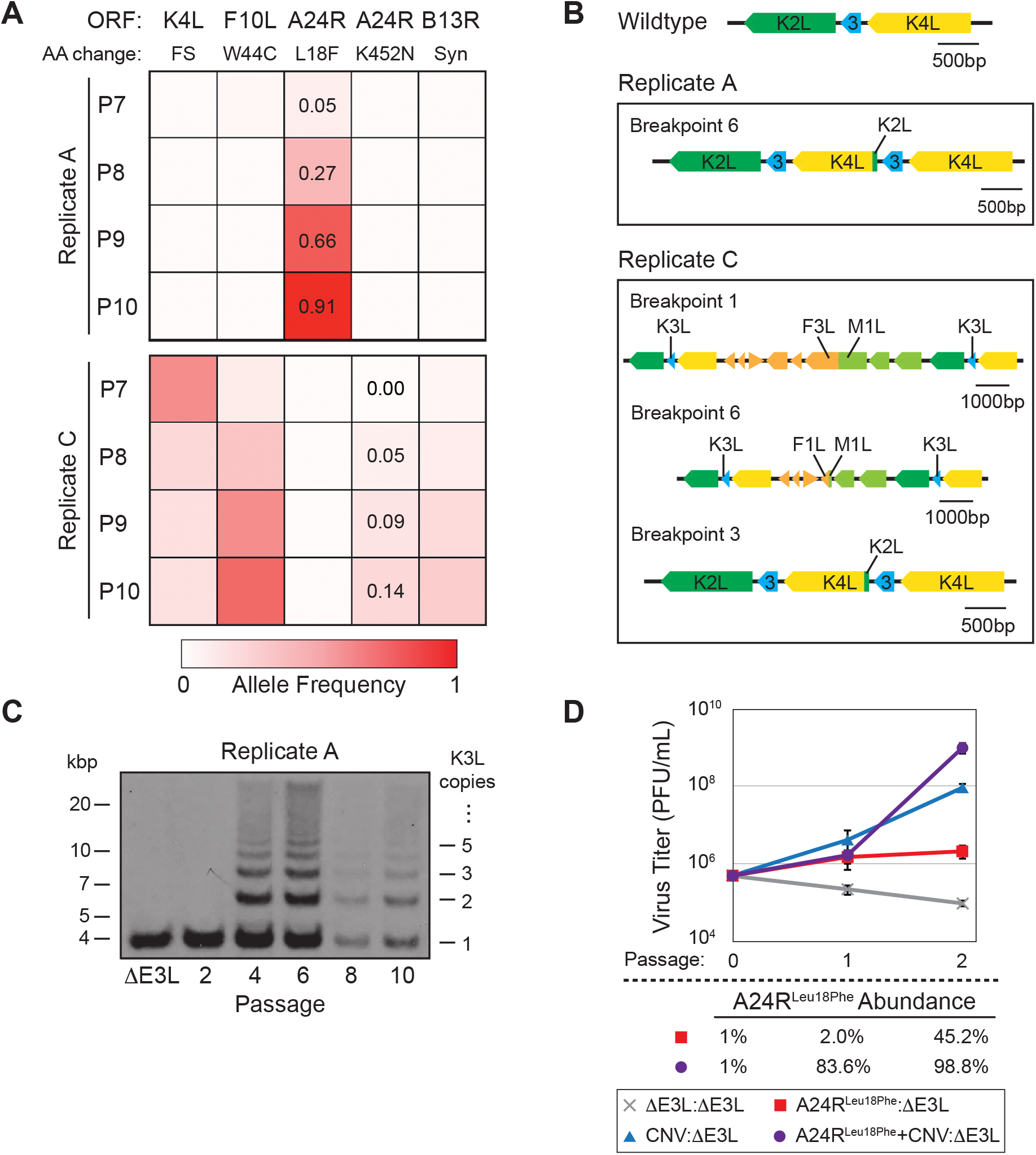
Copy number variation enhances the sweep of a point mutation. (A) Allele frequencies >0.01 from replicates A and C were obtained by deep sequencing of P7–10 virus populations, and position in an ORF and resulting amino acid change are listed above as in Figure 2A. FS = frameshift, Syn = synonymous. (B) Genome structure from direct sequencing across the CNV breakpoints identified by Wham (Table 2) in replicate A and C passage 10 populations. (C) Southern blot was performed on digested viral DNA from indicated viral populations, and labeled with a K3L-specific probe. Size in kbp (left) and number of K3L copies (right) are shown. (D) Viruses were mixed at a ratio of 1:100 as listed, and passaged twice for 48 hours in 1°HF cells (MOI 0.1). Titers were measured in BHK cells by 72-hour plaque assay, as mean PFU/ mL ± standard deviation. A24R Leu18Phe mutation abundance is listed below as input (P0), or as measured by deep sequencing (P1–2).

Increased copy number of the K3L gene in the P10 population provides a clue for how the A24R mutation might rapidly sweep to fixation despite increasing PKR activation. There is only one recombination breakpoint at P10 in replicate A, as opposed to multiple breakpoints in replicates B and C (Figure 5B, Table 2). This suggests that when the selective sweep of the A24R Leu18Phe variant occurred, these viruses also contained a single K3L CNV breakpoint. To determine whether the K3L gene copy number amplification or the A24R Leu18Phe mutation arose first in the virus population, we used Southern blot analysis to measure K3L CNV during experimental evolution of replicate A. We first detected K3L CNV at passage 4, and the proportion of viruses with the population harboring CNV seems to remain steady through passage 10, as indicated by the consistent intensity between bands within each lane (Figure 5C). Since the A24R Leu18Phe substitution did not emerge until passage 7, the earlier appearance of K3L CNV might have preemptively blocked PKR activation that would otherwise be induced by A24R Leu18Phe, facilitating its rapid fixation. Indeed, in cells infected with replicate A P10 virus, which contains both the A24R Leu18Phe substitution and K3L CNV, there is a marked reduction in PKR activation (Figure 3D). This result suggests that K3L CNV can compensate for the activation of PKR induced by the A24R Leu18Phe variant.

To test whether rapid accumulation of the A24R Leu18Phe variant is influenced by K3L copy number, we wanted to track increases of A24R Leu18Phe in virus populations with and without K3L CNV. To do this we infected cells with viruses containing the A24R Leu18Phe substitution alone (A24R^Leu18Phe^), K3L CNV alone (CNV), or the combination of the two genetic changes (A24R^Leu18Phe^+CNV), each starting at a ratio of 1:100 with parental ΔE3L virus (see Methods for strain details). Compared to ΔE3L alone after two passages in 1°HF, the A24R^Leu18Phe^:ΔE3L virus population replicated ~10-fold better, CNV:ΔE3L ~1000-fold, and A24R^Leu18Phe^+CNV:ΔE3L ~10,000-fold (Figure 5D). These data suggest that while either the A24R Leu18Phe variant or K3L CNV is sufficient for a fitness benefit in 1°HF cells, the combination is additive and may therefore facilitate fixation of the A24R Leu18Phe variant. We next analyzed the abundance of the A24R Leu18Phe substitution in the virus populations, to determine if it accumulated faster in the presence of K3L CNV. Starting at 1% of the population, the mutation reached 98.8% abundance after only two passages when multiple copies of K3L were present, compared with 45.2% abundance at passage 2 with a single copy of K3L (Figure 5D). The increased accumulation of the variant in the presence of CNV is consistent with the added fitness benefit for viruses carrying both genetic changes. Thus the A24R Leu18Phe substitution accumulated markedly faster in the presence of K3L CNV, and suggests that the reduction in PKR activation provided by increased K3L expression may have facilitated the rapid rise of the distant A24R mutation.

## Discussion

In this study we found a new means of poxvirus adaptation to innate immune response pathways. Vaccinia has proven a useful model for experimental evolution and continues to reveal the genetic basis of various poxvirus adaptations (14, 23, 35). We charted the rise of multiple point mutations over the course of ten serial infections, which is consistent with adaptive evolution through several independent mechanisms. Most notable among these mutations was the rapid accumulation of Leu18Phe in A24R, which encodes a subunit of the viral RNA polymerase. Unlike in other evolved virus populations, the rise of A24R Leu18Phe appeared in the near absence of other point mutations (Figure 5A), an observation reminiscent of clonal interference, where strongly adaptive point mutations on separate genomes can compete and transiently dominate within asexual populations (41, 42). However, a case of clonal interference seemed unlikely given high rates of recombination in poxviruses (43–46) and the modest replicative advantage we measured in a recombinant strain containing only the A24R Leu18Phe mutation (Figure 3A). We therefore considered the impact of K3L gene copy number amplification as a facilitating event for the rapid accumulation of the A24R Leu18Phe variant during this course of experimental evolution.

We previously found that genomes harboring multiple copies of K3L produced more K3 protein, which resulted in fitness gains for viruses under selective pressure to overcome the antiviral factor PKR (14). In two populations evolved in HeLa cells we observed the additional emergence of a beneficial point mutation in K3L during serial infections, which we predict is more likely to occur in viruses with multiple copies of the gene (14). In this study we observed a beneficial point mutation in a gene distant from the expanded K3L locus (>100,000bp apart) that may benefit from the presence of virus genomes harboring multiple copies of K3L. We speculate that the A24R Leu18Phe substitution alters virus transcription to promote virus replication, but also activates PKR as shown in Figure 3D. However, in the presence of multiple copies of K3L that block PKR activation, the A24R Leu18Phe mutation can rapidly sweep to fixation in virus populations. Consistent with this idea, we observed rapid fixation of the A24R Leu18Phe variant following K3L copy number amplification (Figure 5A, C) and a two-fold increase in the accumulation of A24R Leu18Phe in the presence of multiple K3L copies compared to populations with a single copy of K3L (Figure 5D). These data suggest that adaptive copy number variation can facilitate the rapid accumulation of otherwise modestly beneficial mutations like A24R Leu18Phe. In this way recombination-based CNV could enhance the viability of an expanded set of beneficial mutations that otherwise suffer from tradeoffs (e.g., activation of PKR) and might otherwise be unable to sweep through populations. Given that copy number amplification events are likely transient (14), this foothold may be temporary, as suggested by previous work describing the accumulation of beneficial point mutations causing the collapse of CNV (17). In the current study, CNV of the K3L locus persisted through passage 10 (Figure 5C), but might collapse to a single copy after further rounds of infection. In any case, copy number variation provides an opportunity for mutations to rapidly sweep in genes both undergoing CNV and distant from CNV, despite small initial fitness gains relative to other mutations in the population. Given long periods of evolutionary time, the presence of seemingly simple, beneficial point mutations in virus populations may belie a more volatile history of fixation involving the aid of adaptive, yet transient, gene copy number amplification events.

## Acknowledgements

Author contributions: Experiments were designed by KRC and NCE. Experiments were performed by KRC. Data was analyzed by KRC and ZNK. Analysis tools were contributed by ZNK and MY. The paper was written by KRC and NCE.

We thank Adam Geballe (Fred Hutchinson Cancer Research Center) and Bertram Jacobs (University of Arizona) for reagents. We also thank Adam Geballe for valuable insights and critical assessment of the manuscript. We thank the High-Throughput Genomics Core (University of Utah) for technical assistance.

## Funding

This study was supported by NIH grants R01GM114514 (N.C.E.), R01GM104390 (M.Y.), T32AI055434 (K.R.C.), and T32GM07464 (Z.N.K.). N.C.E. was supported by the Pew Biomedical Scholars program, and the Mario R. Capecchi Endowed Chair in Genetics. The experimental design, data collection and analysis, tools, and decision to submit for publication are our own, and independent of the National Institutes of Health.

